# Imaging of porphyrin-specific fluorescence in pathogenic bacteria *in vitro* using a wearable, hands-free system

**DOI:** 10.1101/2024.05.20.595019

**Authors:** Junhong Sun, Sangeevan Vellappan, Johnathan Akdemir, Liviu Steier, Richard E. Feinbloom, Srujana S. Yadavalli

## Abstract

**Highlights:**

- Fluorescence imaging detects porphyrins in bacteria via natural fluorescence.
- A lightweight, hands-free device enables rapid, non-invasive clinical assessments.
- The study tested 15 bacterial and 2 fungal strains for porphyrin-based autofluorescence.
- 14 bacteria fluoresced on Porphyrin Test Agar, 9 on blood agar plates.
- The REVEAL system aids in diagnosing infections and guiding real-time treatments.

Fluorescence imaging is an effective method for detecting porphyrin production in bacteria, leveraging the natural fluorescence properties of porphyrins. Here we use a simple, lightweight, hands-free device for rapid, non-invasive assessments in clinical settings, microbial research, and diagnostic applications. Specifically in this study, we examined 15 bacterial and 2 fungal strains commonly associated with skin, oral, and/or multi-site infections at wound sites for their ability to autofluoresce based on their porphyrin production. We utilized Remel Porphyrin Test Agar and blood agar plates to monitor red fluorescence over several days of growth under aerobic or anaerobic conditions using the wearable REVEAL FC imaging system with a 405 nm violet excitation headlight paired with eyewear carrying 430 nm emission lenses. Fourteen of the fifteen bacteria produced red fluorescence when grown on Porphyrin Test Agar and nine of the fifteen bacteria also displayed red fluorescence on blood agar plates, consistent with their ability to synthesize porphyrins. Taken together, our results elucidate the sensitivity, effectiveness, and convenience of using wearable technology to detect pathogens that produce porphyrin-specific fluorescence. Consequently, the REVEAL system has immense potential to help diagnose wound infections, direct clinical procedures, and guide treatment options in real-time using fluorescence imaging all while minimizing the risk of contamination.

## 1. Introduction

Microbial infections pose significant threats, especially in wound healing (1, 2). If ignored or not treated promptly, the infection may advance from initial contamination to colonization, local infection, and even systemic infection, potentially leading to severe conditions like sepsis and multiple organ dysfunction syndrome (2). In cases such as the infection in diabetic foot ulcers, timely detection is crucial to prevent potential complications like amputation (3). Similarly, disregarding the initial stages of dental plaque accumulation in the oral cavity can trigger oral health complications, such as gingivitis, periodontitis, dental caries, and even tooth loss (4). Notably, approximately 47.2% of individuals aged 30 years and older are affected by some form of periodontal disease (4).

Considering the rapid spread of infections, when possible, timely detection and effective management are crucial to prevent infections from progressing to severe stages and reduce the risk of complications, such as tissue necrosis and the need for amputation (5, 6). Traditional methods of detecting and confirming bacterial infections, such as culturing and susceptibility testing, are laborious and time-consuming, taking days to yield results (7). In recent years, there has been a development of fluorescence-based approaches for real-time visualization of intrinsic fluorescence from porphyrins produced in bacteria in wound sites and oral plaques, enabling earlier detection (8–11). Fluorescence-based detection of bacteria relies on the production of porphyrins, a crucial metabolic intermediate essential for heme biosynthesis and various biological processes (12). Heme, synthesized by pathogens or obtained from host sources during infection, plays a pivotal role in energy generation through the electron transport chain (12). Heme biosynthesis is vital to both aerobic and anaerobic respiration in prokaryotes (13). In laboratory settings, heme precursors can be provided to microorganisms through specialized media like porphyrin test agar, or blood agar enriched with 5% sheep’s blood cells (14, 15). As pathogens synthesize porphyrins, exposure to violet light (∼400-450 nm) leads to their excitation, emitting light at distinct wavelengths within the red fluorescence at 620-630 nm (16, 17). Like porphyrin, pyoverdines are siderophores uniquely produced by *Pseudomonas spp.* for iron acquisition, which can be visualized in cyan spectral bands (18). This fluorescence emission facilitates identifying and visualizing pathogens, thereby enhancing diagnostic procedures.

Currently, the real-time fluorescence-guided detection of pathogens uses a handheld device (8). Serena and colleagues suggest that the use of fluorescence imaging has resulted in changes to 69% of the treatment plans and most importantly, there has been a significant improvement in patient care in 90% of study wounds (19). This comes as a great benefit to the patient and the clinician. However, as this technology requires a clinician to use both hands, it hampers the clinician’s ability to provide treatment during real-time fluorescence visualization. The adoption cost of this technology requires capital expenditure and incurs ongoing operational costs that limit adoption. Hence, there is a need to develop alternative tools to the existing non-wearable methods, which can allow for rapid and non-invasive assessment.

In this study, we report on a new, wearable/ hands-free fluorescence visualization technology (REVEAL FC wearable imaging system, Design for Vision Inc., Fig 1a) that will enable clinicians to evaluate the infection at the wound site based on real-time visualization of pathogens and simultaneously make active treatment decisions. Specifically, we sought to test and characterize porphyrin detection *via* autofluorescence *in vitro* for some of the common pathogens involved in skin, oral, and other (respiratory, intestinal, urinary tracts, or blood) infections over a period of multiple days. We included 15 bacteria and 2 fungi in our analysis. Our results indicate the ability of REVEAL FC to visualize porphyrin-specific red-orange or pink fluorescence in 14 of the 17 strains tested on porphyrin test agar (Fig 1b). Moreover, we also observed a fluorescence signal for 9 of the 17 strains grown on blood agar plates, suggesting a strong potential for the detection of microbial infections in wound sites in a clinical setting using the REVEAL FC wearable technology.

**Figure 1.**
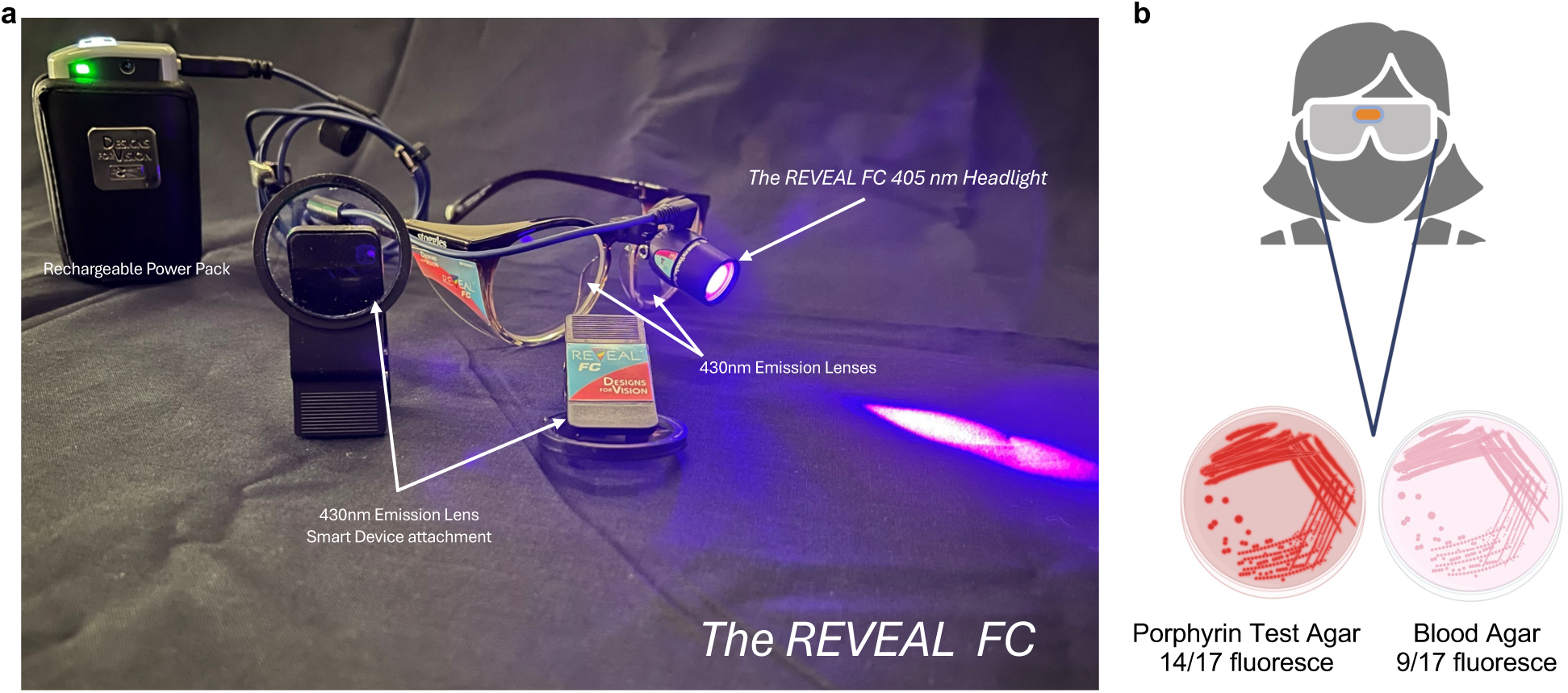
Overview of porphyrin-specific fluorescence detection in pathogenic microbes using the REVEAL FC imaging system. a) The REVEAL FC is a wearable imaging system composed of a headlight that provides excitation at 405 nm and observational eyewear, which includes the emission filter at 430 nm. This system includes a rechargeable power pack and smart device attachment for simple and effective visualization and imaging of microbial samples. b) 17 microbial species (15 bacteria and 2 fungi) commonly associated with skin, oral, and other infections were tested for autofluorescence of porphyrins. 14 of them exhibited fluorescence in shades ranging from red/orange to pink on porphyrin test agar, while 9 of them displayed fluorescence on blood agar.

## 2. Materials and Methods

### 2.1. Strains

*E. coli* K-12 MG1655 strain is from the Yadavalli lab collection. The following strains, *Pseudomonas aeruginosa* PAO1, *Klebsiella pneumoniae* ATCC 13883, *Staphylococcus aureus* USA300_LAC, a community-acquired MRSA strain, and *S. Typhimurium* 14028, were gifts from Drs. Bryce Nickels (Rutgers), Huizhou Fan (Rutgers), Jeffrey Boyd (Rutgers), and Dieter Schifferli (University of Pennsylvania). All other bacterial and fungal strains used in the study were purchased from the American Type Culture Collection (ATCC). See Table S1 for additional details on the bacterial and fungal strains used in this study.

### 2.2. Media and growth conditions

For routine growth, bacterial and fungal strains were cultured in liquid or solid agar medium, either aerobically or anaerobically depending on the organism as specified in Table S1. Agar plates and culture tubes were then incubated at 37°C with aeration (for aerobic growth) or an anaerobic growth chamber (for anaerobic growth), depending on a given strain’s specific growth requirement.

To assess porphyrin production and monitor the associated fluorescence, the strains were streaked out and grown on Remel Porphyrin Test Agar (PTA, Thermo Fisher Scientific), and/or blood agar (BA, tryptic soy agar with 5% sheep blood, Thermo Fisher Scientific). Note that both PTA and BA plates contain heme precursors. For strains grown aerobically, PTA and BA plates were incubated for 5 days at 37°C. Plates were pulled out for imaging daily and returned to incubation until day 5. For anaerobic strains, PTA and BA plates were incubated for 10 days in the anaerobic growth chamber, including one plate per strain for each time point (days 2, 4, 7, and 10) to be pulled out for imaging. At the indicated time points, plates were imaged using the REVEAL FC technology as described below.

### 2.3. Fluorescence imaging

For strains grown aerobically, imaging was performed daily from day 1 to 5. For strains grown in the anaerobic chamber, one plate corresponding to each strain was removed from the chamber on days 2, 4, 7, and 10, and imaged. Fluorescence visualization and imaging were performed using the REVEAL FC, a wearable system equipped with the following: a 405 nm headlight for excitation, an observational eyewear with an emission filter at 430 nm, a rechargeable power pack (2 included) with 9 hours of runtime per pack, a smart device attachment for simple and easy documentation. For this study, a smartphone adapter with a 430 nm emission filter was attached to an Apple iPhone 12 to capture the images. The images were taken in a dark room. Images were analyzed and processed using ImageJ (20).

## 3. Results

### 3.1. REVEAL FC imaging system for *in vitro* visualization and imaging of microbial fluorescence

Fluorescence visualization requires an excitation source and an emission filter to eliminate the excitation illumination to maximize the observation of the fluorescent response. The REVEAL FC developed by Designs for Vision Inc. is a wearable system that is composed of a headlight that provides excitation and observational eyewear that includes the emission filter (Fig 1). It is well documented that porphyrins fluoresce when exposed to light in the 400-409nm range (21).

Porphyrins are produced as by-products by several microorganisms, including pathogens of interest. The REVEAL FC headlight utilizes an LED at 405 nm wavelength. Even though the peak of this LED is at 405nm, well within the excitation range of porphyrins, LEDs are relatively unfocused. The LED has a wide spectral bandwidth with a transmission range (∼380 – 450 nm) that is too broad for specific excitation of porphyrins (Fig S1) and the spot is too wide to respond to lensing. The REVEAL FC system uses proprietary and patented technology involving short wave pass edge filters (shortpass filters) to narrow the spectral bandwidth. Focusing the energy of the LED to a fixed spot provides uniform illumination within the wavelengths that strongly excite porphyrins (400 – 425 nm) while eliminating unnecessary wavelengths that could inhibit the visualization of the fluorescence response. The emission filter is incorporated into a pair of glasses, which removes all wavelengths less than 430 nm providing a clear image of the field with no tinting. All the excitation light is removed by the emission filter observational glasses allowing maximum visualization of the fluorescence response. For documentation, a smartphone adapter is provided that contains the same emission filter (Fig 1). This is intended to be attached to any smart device to document and capture the fluorescent images.

### 3.2. Porphyrin-specific fluorescence in bacteria and fungi causing skin infection

We explored the fluorescent characteristics of seven prevalent bacterial species and two fungal species known for causing skin infections or thriving on open skin wounds using the REVEAL FC system. Among the bacteria tested, three are Gram-positive (*S. aureus*, *C. minutissimum*, *C. acnes*) while the remaining four, *A. hydrophila*, *S. marcescens*, *V. vulnificus* and *P.aeruginosa* are Gram-negative (Table 1). These bacterial strains were grown aerobically except for *C. acnes*, which is an obligate anaerobe. All of them demonstrate red fluorescence to different extents as monitored over several days and described below (Table 1, Fig 2).

**Figure 2.**
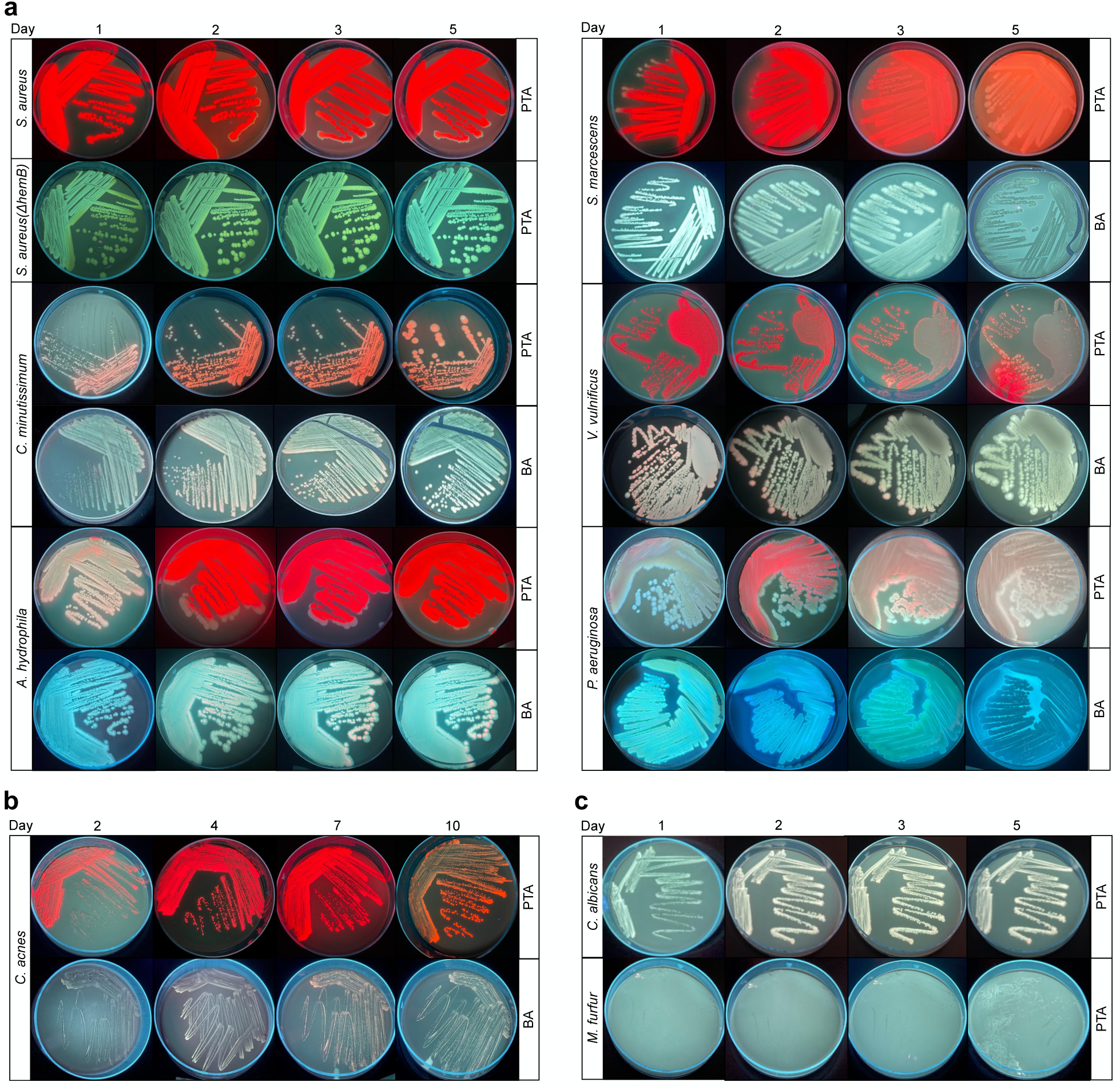
Porphyrin-specific fluorescence detection in microbes associated with skin infection. a) Fluorescence-based imaging of bacteria on porphyrin test agar (PTA) and/or blood agar under aerobic conditions after 1, 2, 3, and 5 days of growth. Bacterial strains *Staphylococcus aureus* and its mutant *ΔhemB*, *Corynebacterium minutissimum*, *Aeromonas hydrophila*, *Serratia marcescens*, *Vibrio vulnificus*, and *Pseudomonas aeruginosa* are labeled as *S. aureus*, *S. aureus ΔhemB*, *C. minutissimum*, *A. hydrophila*, *S. marcescens*, *V. vulnificus*, and *P. aeruginosa*, respectively. b) Fluorescence-based imaging of bacteria grown on porphyrin test agar (PTA) and/or blood agar under anaerobic conditions after 2, 4, 7, and 10 days of growth. The bacterial strain *Cutibacterium acnes* is labeled as *C. acnes*. c) Fluorescence-based imaging of fungi on porphyrin test agar (PTA) and/or blood agar under aerobic conditions after 1, 2, 3, and 5 days of growth. Fungal strains *Candida albicans* and *Malassezia furfur* are labeled as *C. albicans* and *M. furfur*, respectively. Images are representative of four biological replicates.

**Table 1.** List of bacterial and fungal strains plated on Porphyrin Test Agar (PTA) or blood agar (BA) and tested for fluorescence production.

| Strain | Category | Growth | Incubation<br>time in<br>days | Fluorescence |  |
| --- | --- | --- | --- | --- | --- |
|  |  |  |  | PTA | BA |
| Gram-positive bacteria |  |  |  |  |  |
| *Corynebacterium*<br>*minutissimum* | skin | aerobic | 5 | Red/orange | Pink |
| *Cutibacterium*<br>*acnes* (previously<br>*Propionibacterium*<br>*acnes*) | skin | anaerobic | 10 | Red | Red |
| *Staphylococcus*<br>*aureus* | skin | aerobic | 5 | Red | – |
| *Staphylococcus*<br>*aureus* (ΔhemB) | | aerobic | 5 | – | – |
| *Streptococcus*<br>*mutans* | oral | aerobic | 5 | – | – |
|  |  | anaerobic | 10 | – | – |
| *Listeria*<br>*monocytogenes* | other<br>(blood) | aerobic | 5 | Red | – |
| *Streptococcus*<br>*pyogenes* | other<br>(respiratory<br>tract) | anaerobic | 10 | Red | Red |
|  |  | aerobic | 5 | – | – |
| Gram-negative bacteria |  |  |  |  |  |
| *Aeromonas hydrophila* | skin | aerobic | 5 | Red | Pink |
| *Pseudomonas aeruginosa* | skin, multi-site | aerobic | 5 | Red | Cyan |
| *Serratia marcescens* | skin, urinary tract | aerobic | 5 | Red | Pink |
| *Vibrio vulnificus* | skin | aerobic | 5 | Red | Pink |
| *Porphyromonas gingivalis* | oral | anaerobic | 10 | Red | Red/black |
| *Escherichia coli* | other (intestinal, urinary tract) | aerobic | 5 | Red | – |
| *Klebsiella pneumoniae* | other (multi-site) | aerobic | 5 | Red | – |
| *Proteus mirabilis* | other (urinary tract) | aerobic | 5 | Pink | Cyan |
| *Salmonella typhimurium* | other (intestinal tract) | aerobic | 5 | Red | – |
| **Fungi** |  |  |  |  |  |
| *Candida albicans* | skin | aerobic | 10 | – | – |
| *Malassezia furfur* | skin | aerobic | 10 | – | – |

In aerobic growth, all bacterial species exhibited fluorescence, while the fungi did not (Table 1, Fig 2a). *Staphylococcus aureus* displayed intense red fluorescence on porphyrin test agar (PTA) from day 1, maintaining its intensity through day 5. No fluorescence was observed on blood agar (BA), whose base is tryptic soy medium (Table S1) that is known to exhibit background fluorescence that can mask the porphyrin-specific signal (8). Through the Boyd lab at Rutgers, we had access to *S. aureus hemB* mutant strain, which is deficient in heme acquisition and porphyrin production (22). Consistently, *S. aureus hemB* mutant showed no fluorescence throughout the five days, serving as a negative control (8). *Serratia marcescens* also showed fluorescence on both PTA where the red fluorescence was visible from day 1, intensifying with each passing day until day 5. On BA, *S. marcescens* seemed to show a faint pink fluorescence on day 1, though there was no signal on days 2 and 5. *Corynebacterium minutissimum* exhibited red fluorescence on PTA starting from day 1, with increased intensity on days 2 and 3, persisting at least until day 5 when all colonies exhibited red fluorescence.

Interestingly, we detected red fluorescence for *C. minutissimum* on BA from day 1, reaching maximum intensity by days 2-3, which indicates a signal higher than the background for BA and the fluorescence diminished by day 5. Similarly, for *Vibrio vulnificus*, bright red fluorescence was visible on days 1 and 2, gradually diminishing by days 3 and 5. On BA, *V. vulnificus* showed red fluorescence as observed on days 1 and 2, with the intensity decreasing and nearly disappearing by day 5. *Aeromonas hydrophila* displayed red fluorescence on PTA from day 1, intensifying remarkably by day 2 and the bright red signal stayed steady until day 5. Similar to *C. minutissimum* and *V. vunificus*, we observed red fluorescence for *A. hydrophila* on BA but with a delayed appearance on day 2, progressively intensifying by day 5. *Pseudomonas aeruginosa* exhibited red fluorescence on PTA, with a weak signal on day 1, transitioning to a brighter red signal by day 2 and continuing into day 5 (Fig 2a). On BA, *P. aeruginosa* exhibited cyan fluorescence on day 1, persisting until day 5. *Pseudomonas spp.* are known to synthesize a variety of phenazine pigments including pyoverdines (23), which contribute to the observed cyan fluorescence in our *P. aeruginosa* PAO1 strain.

The fluorescence changes in anaerobes were observed over a longer period, extending up to 10 days, due to their slower growth rate. *Cutibacterium acnes*, an aerotolerant anaerobe (24), exhibited fluorescence on both PTA and BA (Fig 2b). It displayed red fluorescence from day 2, intensifying by day 4 and reaching peak intensity around day 7, but the fluorescence intensity reduced on day 10. On BA, very faint red fluorescence was observed on day 2, which is more evident on day 4, intensifying by day 7, and diminished by day 10. Fungal strains *Candida albicans* and *Malassezia furfur* did not show red fluorescence, which is consistent with previous studies (8) (Fig 2c).

### 3.3. Porphyrin-specific fluorescence in bacteria causing oral infection

We examined two bacterial pathogens known for causing oral infections: *Streptococcus mutans*, a Gram-positive bacterium typically inhabiting dental plaque (25), and *Porphyromonas gingivalis*, a Gram-negative obligate anaerobe associated with pulpal infections, oral abscesses, and periodontitis (26). *S. mutans* showed no fluorescence under aerobic growth but appeared to show a faint orange signal under anaerobic conditions (Table 1, Fig 3). The second oral pathogen, *P. gingivalis* formed red colonies on PTA, beginning as early as day 2 and 4, but diminishing by day 7 and becoming less visible by day 10 (Fig 3b). On BA, it exhibited no fluorescence on day 2 but developed red fluorescence by day 4. By day 7 on BA, there is a reduction in the red fluorescence and *P. gingivalis* starts to show black pigmentation, which is more prominent by day 10 (Fig 3b). The black-pigmented colonies on blood agar are associated with the accumulation of heme complexes as noted previously (27).

**Figure 3.**
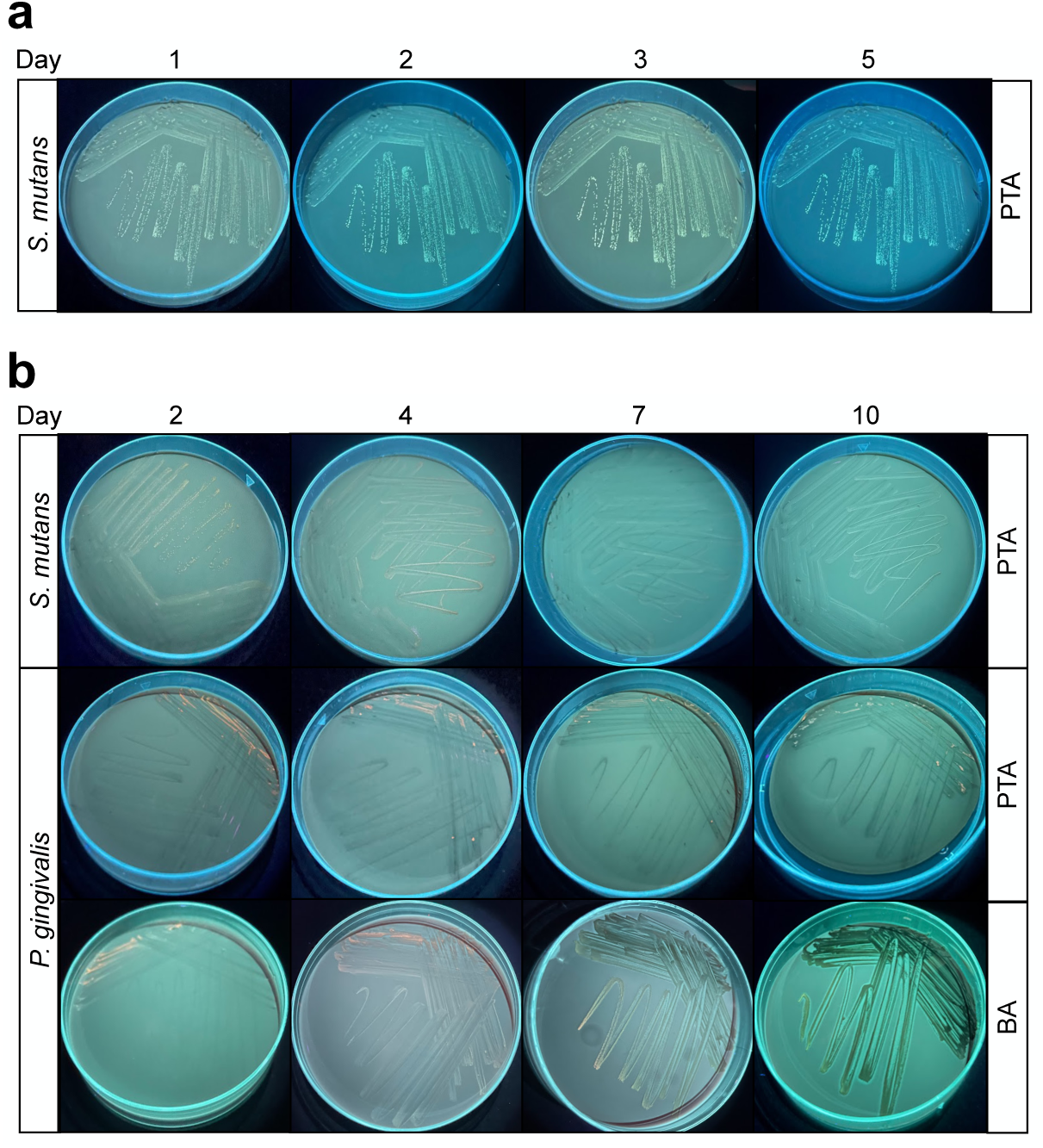
Porphyrin-specific fluorescence detection in bacteria associated with oral infection. a) Fluorescence-based imaging of bacteria on porphyrin test agar (PTA) and/or blood agar under aerobic conditions after 1, 2, 3, and 5 days of growth. The bacterial strain *Streptococcus mutans* is labeled as *S. mutans*. b) Fluorescence-based imaging of bacteria grown on porphyrin test agar (PTA) and/or blood agar under aerobic conditions after 2, 4, 7, and 10 days of growth. Bacterial strains *Streptococcus mutans* and *Porphyromonas gingivalis* are labeled as *S. mutans* and *P. gingivalis*, respectively. Images are representative of four biological replicates.

### 3.4. Porphyrin-specific fluorescence in bacteria causing multi-site/other infections

In this section, we tested six bacterial strains that exhibit a diverse range of pathogenic behaviors across distinct anatomical sites and not being exclusively confined to cutaneous or oral infection (Table 1). Two of the six bacterial strains are Gram-positive, while the remaining four are Gram-negative. Five of these species (*Listeria monocytogenes*, *Escherichia coli*, *Klebsiella pneumoniae*, *Salmonella typhimurium,* and *Proteus mirabilis*) demonstrate fluorescence under aerobic conditions, whereas one (*Streptococcus pyogenes*) exhibits fluorescence exclusively under anaerobic conditions (Fig 4).

**Figure 4.**
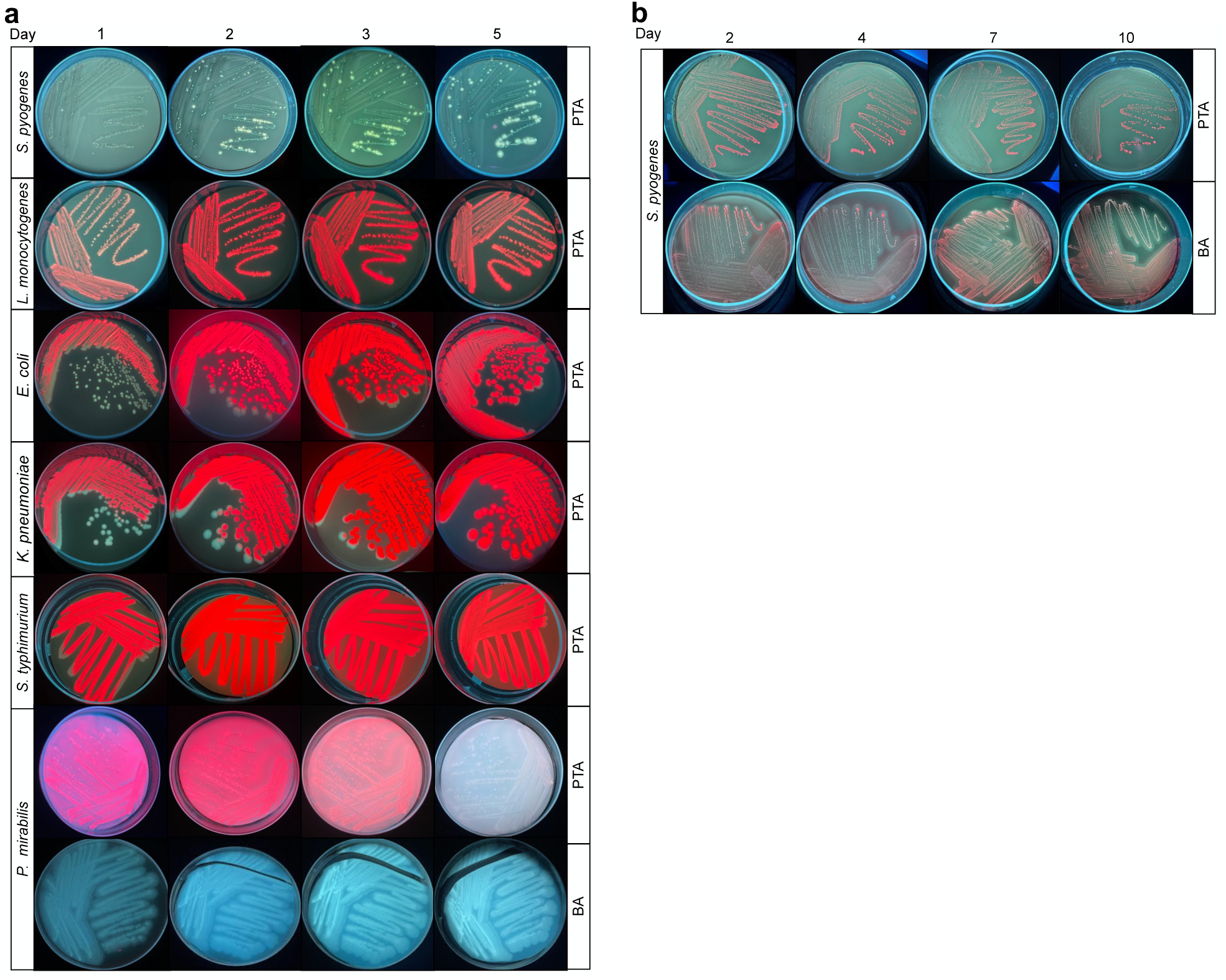
Porphyrin-specific fluorescence detection in bacteria associated with other infections. a) Fluorescence-based imaging of bacteria on porphyrin test agar (PTA) and/or blood agar under aerobic conditions after 1, 2, 3, and 5 days of growth. Bacterial strains *Streptococcus pyogenes*, *Listeria monocytogenes*, *Escherichia coli*, *Klebsiella pneumoniae*, *Salmonella typhimurium*, and *Proteus mirabilis* are labeled as *S. pyogenes*, *L. monocytogenes*, *E. coli*, *K. pneumoniae*, *S. typhimurium*, and *P. mirabilis*, respectively. b) Fluorescence-based imaging of bacteria grown on porphyrin test agar (PTA) and/or blood agar under aerobic conditions after 2, 4, 7, and 10 days of growth. The bacterial strain *Streptococcus pyogenes* is labeled as *S. pyogenes*. Images are representative of four biological replicates.

Under aerobic growth conditions, *S. pyogenes*, a facultative anaerobe (28), did not exhibit any fluorescence on PTA (Fig 4a). However, when *S. pyogenes* was grown under anaerobic growth conditions, red fluorescence was readily visible from day 2 and persisted through days 4, 7, and 10 on both PTA and BA (Fig 4b). *L. monocytogenes* exhibited fluorescence on PTA from day 1, intensifying by day 2 with a slight reduction in intensity by day 5 (Fig 4a). Three strains, *E. coli*, *K. pneumoniae*, and *S. typhimurium* displayed red fluorescence from day 1 with an increased intensity on day 2, which remained strong and stable through day 5. Intriguingly, *P. mirabilis* exhibited bright pink fluorescence on PTA on day 1, intensifying on days 2 and 3, but reducing in intensity by day 5. On BA, *P. mirabilis* showed cyan fluorescence, which was visible on day 1, gradually intensifying by day 3 and remaining consistent thereafter. Secondary metabolites such as phenazines are thought to be produced by a variety of bacteria (29) and the cyan fluorescence observed here for *P. mirabilis* grown on BA may be due to the accumulation of a pyoverdine-like pigment.

## 4. Discussion and conclusions

Wearable technology in clinical practice and diagnostics encompasses devices such as smartwatches, fitness trackers, smart clothing, and implantable sensors, which collect data on various physiological parameters such as heart rate, activity levels, sleep patterns, blood glucose levels, and more (30, 31). These devices have a broad scope of applications in monitoring chronic conditions, fitness, wellness, rehabilitation, clinical trials, and mental health as well as early diagnosis and prevention. Additionally, there has been progress in developing wearable devices for clinicians and healthcare professionals ranging from smartwatches and augmented reality glasses to biometric sensors to enhance diagnosis, surgical planning, precision, and hands-free communication (32, 33).

In this study, we use the REVEAL FC imaging system developed by Designs for Vision Inc., a lightweight and comfortable hands-free device that can be customized to the operator’s ocular specifications (Fig 1, S1). It minimizes cross-contamination risks and enhances objective assessment capabilities, overcoming limitations posed by bulky and tedious traditional tools. Here we systematically analyzed 15 bacterial and 2 fungal pathogens (Table 1, S1) for red fluorescence associated with porphyrin production *in vitro* under aerobic (days 1 through 5) or anaerobic (days 2 through 10) growth conditions. By plating strains on porphyrin test agar (PTA), we detected a strong red fluorescent signal in all but one (*S. mutans*) of the bacterial strains tested. *S. pyogenes* displayed red fluorescence but only under anaerobic growth conditions. Two other *Streptococcus* strains do not display red fluorescence as documented previously (8), and some bacteria from this genus are known to be deficient in heme biosynthesis (12, 34). Remarkably, red fluorescence has been reported for *S. mutans* in a dentin caries model pointing to a potential media- and context-dependent mechanism for porphyrin production in this bacterium (35).

For a majority of the bacterial strains, the signal was evident by day 1 and in a few cases, depending on the individual bacterial strain, there was a delayed onset of fluorescence. We also plated the strains on blood agar and found that 9 of the 15 bacteria displayed red fluorescence.

It is worth noting that the trypic soy blood agar medium has background fluorescence that typically masks any signal from the bacteria, however, REVEAL was able to detect the red signal above the threshold of the background, suggesting that this system is highly sensitive. Unsurprisingly, the two fungal strains included in this study, *C. albicans* and *M. furfur* did not show porphyrin-dependent red fluorescence, suggesting that they may not produce heme and instead acquire heme from their environments (36).

This method, like other porphyrin-based approaches, is limited by the fact that not all bacteria produce porphyrins. Internal and external factors can also impact porphyrin production, affecting detection. While the in vitro agar plate model may not fully replicate real-world infection conditions, it provides a foundational step to guide clinicians.

Overall, wearable technology for healthcare professionals enhances medical practice by enabling faster detection and intervention, improving patient care, increasing efficiency, and supporting more informed clinical decision-making. The use of wearable technology is directly linked to improved patient outcomes in clinical settings (37). For instance, the detection of bacteria in the oral cavity plays a crucial role in diagnosing and treating infections, as they serve as indicators for infected tissue and dental plaque (15). Previous work has shown that fluorescence imaging using the REVEAL system can help with diagnosis and treatment guidance in cariology, oral hygiene, and peri-implantitis (35, 38–40).

More broadly, the smart wearable fluorescence imaging system described here has immense potential for diagnosis, treatment, and many other applications across pharmaceutical, healthcare, food, and agricultural industries.

## Acknowledgement

We thank Drs. Meliza Talaue, Xuesong Zhang, and Dr. Martin Blaser’s laboratory at the Center for Advanced Biotechnology and Medicine at Rutgers University for the access and use of an anaerobic chamber.

## Ethics statement

This study did not include human samples. The use of human pathogens in the licensed BSL-2 laboratory was approved by the institutional biosafety committee. All biosafety regulations were followed during the conduct of this study.

## Declaration of generative AI in scientific writing

The current version of ChatGPT (GPT-4o) was used to improve the readability and clarity in the abstract and introductory sections of this paper. The authors declare that generative AI was not used for scientific writing or in creating figures or images included in the manuscript.

## CRediT authorship contribution statement

Junhong Sun: Data curation, Investigation, Validation. Sangeevan Vellappan: Data curation, Visualization, Writing-Original draft preparation. Johnathan Akdemir: Data curation, Visualization. Liviu Steier: Methodology, Resources. Richard E. Feinbloom: Methodology, Resources. Srujana S. Yadavalli: Conceptualization, Funding acquisition, Supervision, Writing-Original draft preparation, Writing-Reviewing and Editing.

## Data Availability Statement

The authors declare that all data supporting the findings of this study are available within the main article and its supplementary information files.

---

**Figure S1.**
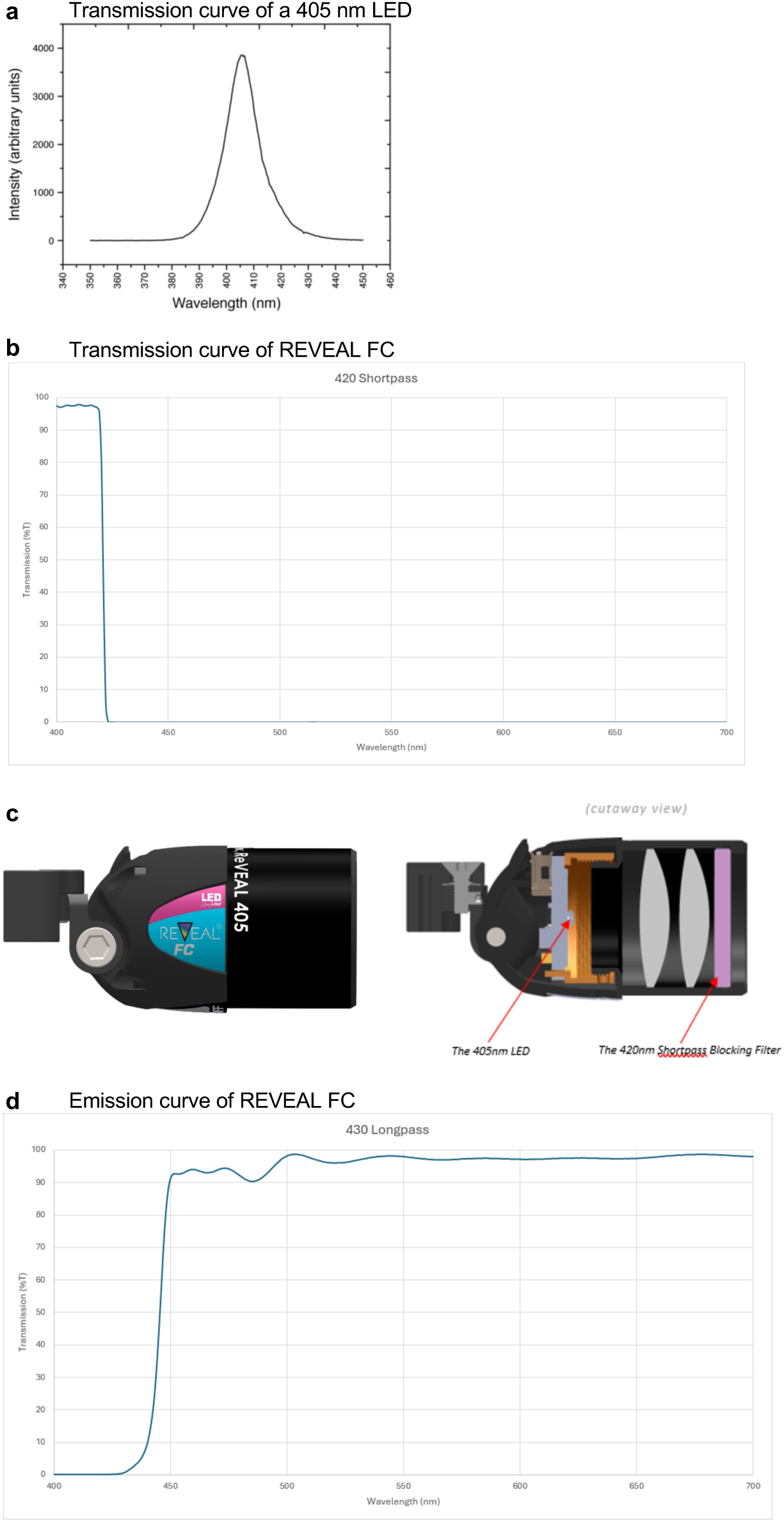
Fluorescence emission spectra for the 405 nm LED. a) Representative transmission curve of a typical 405 nm LED showing light transmission across a range of wavelengths (380 – 450 nm). b) Transmission curve of REVEAL FC using a 420 nm shortpass filter focusing fluorescence excitation at 400 – 425 nm. c) Diagram of the REVEAL FC lens with a cross-sectional depiction of the 405 nm LED and the 420 nm shortpass cut-off filter. d) Emission curve of REVEAL FC lens showing emission at wavelengths ≥ 430 nm.

**Table S1.**
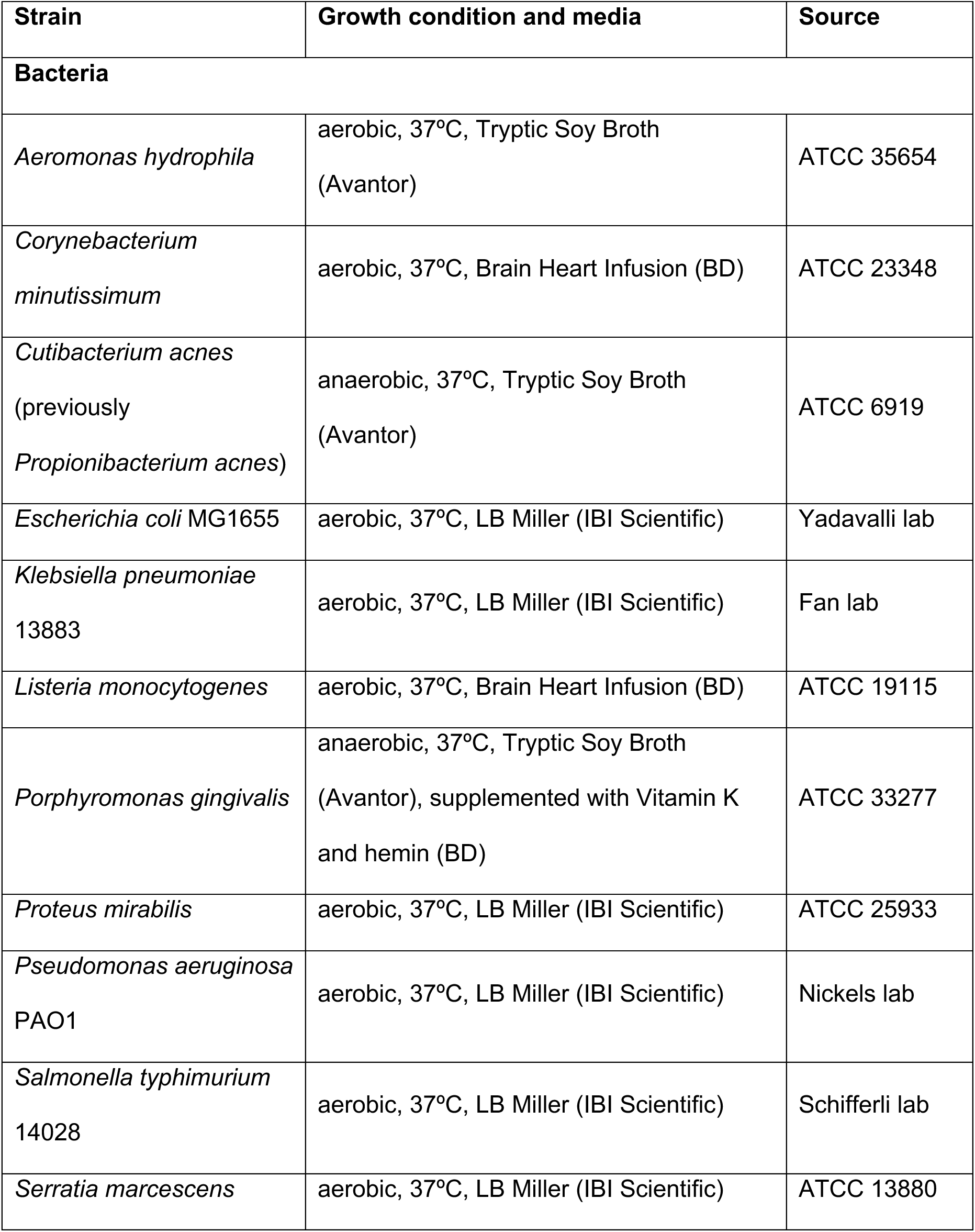
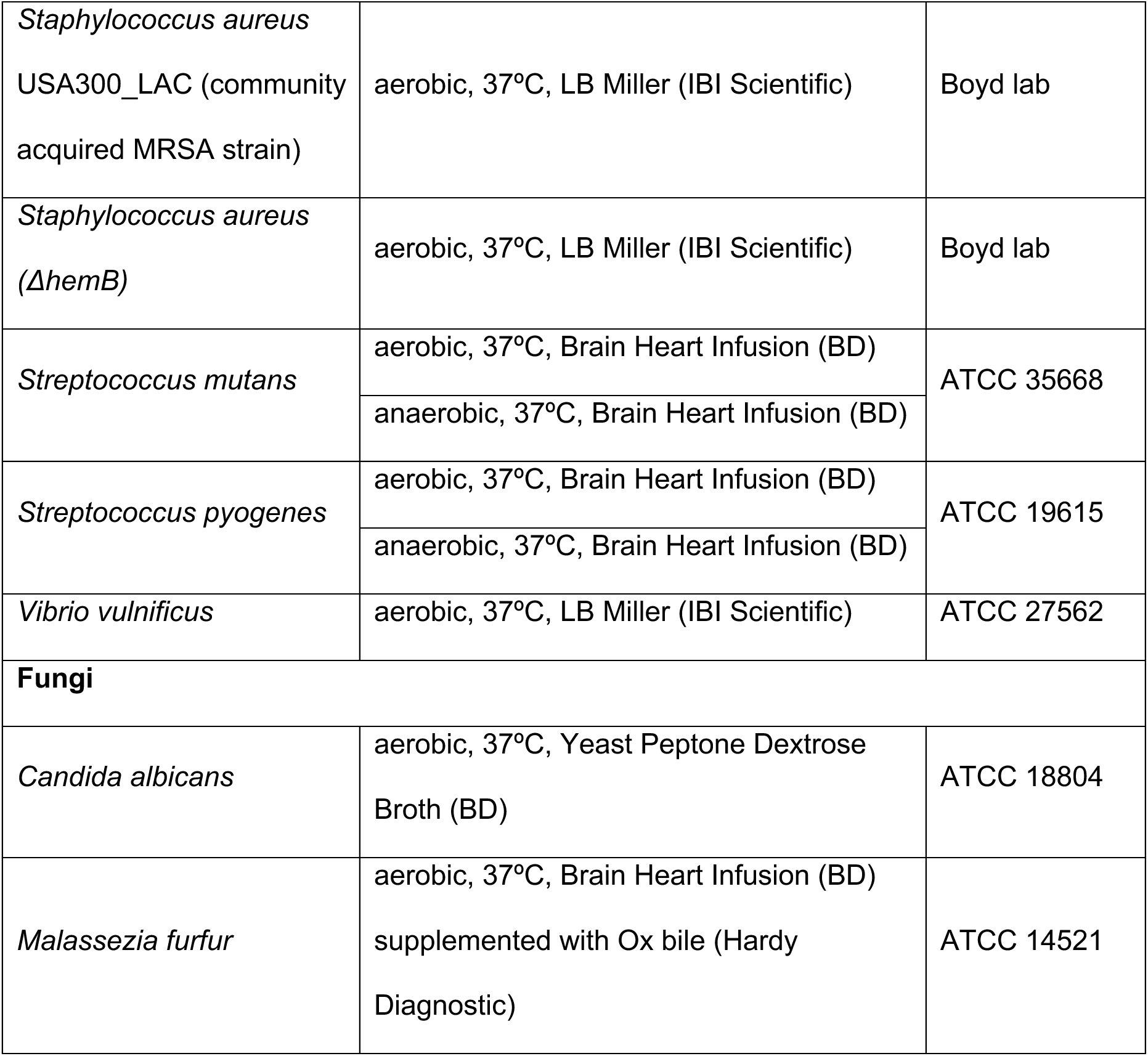
List of strains, growth conditions, and liquid culture media used in the study.

